# X-chromosome upregulation is dynamically linked to the X-inactivation state

**DOI:** 10.1101/2020.07.06.189787

**Authors:** Antonio Lentini, Christos Coucoravas, Nathanael Andrews, Martin Enge, Qiaolin Deng, Björn Reinius

## Abstract

Mammalian X-chromosome dosage balance is regulated by X-chromosome inactivation (XCI) and X-chromosome upregulation (XCU), but the dynamics of XCU as well as the interplay between the two mechanisms remain poorly understood. Here, we mapped XCU throughout early mouse embryonic development at cellular and allelic resolution, revealing sex- and lineage-specific dynamics along key events in X-chromosome regulation. Our data show that XCU is linearly proportional to the degree of XCI, indicating that dosage compensation ensues based on mRNA levels rather than number of active X chromosomes. In line with this, we reveal that the two active X chromosomes in female naïve embryonic stem cells are not hyperactive as previously thought. In all lineages, XCU was underlain by increased transcriptional burst frequencies, providing a mechanistic basis *in vivo*. Together, our results demonstrate unappreciated flexibility of XCU in balancing X-chromosome expression, and we propose a general model for allelic dosage balance, applicable for wider mechanisms of transcriptional regulation.

## Main

In therian mammals, the X chromosome is present as two copies in females and one in males. Correct balance of gene dosage is vital for homeostasis and normal cell function and two major, X-chromosome-wide, mechanisms ensure balanced genomic expression. XCI silences one X allele in females, equalizing expression between the sexes, and XCU increases X-linked expression to resolve dosage imbalance against autosomal gene networks [reviewed in ref. (*1*)]. Although the existence of XCU is well supported by relative measurements between X and autosomes (*2*–*6*), the dynamics of its establishment and maintenance remain elusive (*6*–*8*). Such unraveling would require quantitative gene expression measurements at cellular and allelic resolution, only recently made possible by allele-sensitive single-cell RNA-seq (scRNA-seq), and analytical disentanglement of other allelic processes at cellular level such as stochastic allelic expression and random XCI (rXCI) (*9*). Here, we have mapped the XCU process in mouse at allelic and cellular resolution throughout its establishment *in vitro* and *in vivo*, revealing fundamental features and a surprising plasticity in this process, and we provide a unified model of this chromosome-wide mode of gene regulation.

## Results

### Establishment of X-upregulation during mESC differentiation

We started by culturing naïve male and female mouse embryonic stem cells (mESCs) derived from F1 hybrids of strains C57BL/6J and CAST/EiJ (hereafter C57 and CAST, respectively), and subsequently differentiated female mESCs into epiblast stem cells (EpiSCs) to induce rXCI (**Fig. 1a, Methods**). We harvested naïve mESCs and EpiSCs throughout differentiation (day 1, 2, 4 and 7) and subjected cells to scRNA-seq using the recently published Smart-seq3 method (*10*) (**Fig. 1a, Methods**), allowing sensitive allele- and molecule-level analysis at cellular resolution (**Supplementary Fig. 1a**). Differentiation induced distinct expression changes accompanied by loss of pluripotency factors (*e.g. Sox2* & *Nanog*) and induction of lineage-specific factors (*e.g. Fgf5* & *Krt18*) (**Fig. 1b-c**, **Supplementary Fig. 1b and Supplementary Table 1**) as well as enrichment for associated pathways (**Supplementary Fig. 1c and Supplementary Table 1**). *Xist* induction was detected already by differentiation day 1 (**Supplementary Fig. 1d**) and accompanied by the transition from biallelic to monoallelic X-chromosome expression (**Fig. 1d**), signifying successful rXCI. To investigate XCU, we stratified female cells based on rXCI status into active (XaXa) and inactive (XaXi) states inferred from allelic expression (**Fig. 1d, Methods**) and calculated X-chromosome expression per cell and allele. Intriguingly, we detected chromosome-wide upregulation of the active X allele in female cells with established rXCI and in male mESCs, but lack thereof in female cells of XaXa state (**Fig. 1e-f**). We calculated total expression of the different X states revealing that the combined output of two moderately active XaXa alleles was higher than that of a single hyperactive allele (FDR_TPM_ ≤ 4.35 × 10^−34^, FDR_UMI_ ≤ 6.86 × 10^−5^, FDR corrected Mann-Whitney *U* test; **Fig. 1g-h and Supplementary Fig. 1e**). Furthermore, gene-wise fold-changes of the active allele transitioning from XaXa into XaXi state (**Fig. 1i**, shown for both C57 and CAST alleles) demonstrated distinct XCU, with fold-changes below two (median X_C57_ = 1.62, P_C57_ = 1.41 × 10^−40^ and median X_CAST_ = 1.67, P_CAST_ = 5.72 × 10^−48^, Wilcoxon signed-rank test). These findings are important as they alter the notion that XCI acts on an hyperactive XaXa state (*3*, *5*, *11*). It furthermore directly explains why measured female-to-male X-chromosome expression fold-change in naïve ESC tend to be less than two-fold (*5*, *11*), and is consistent with the notion that expression dosage of two active X alleles blocks or delays differentiation of mESCs (*12*).

**Figure 1.**
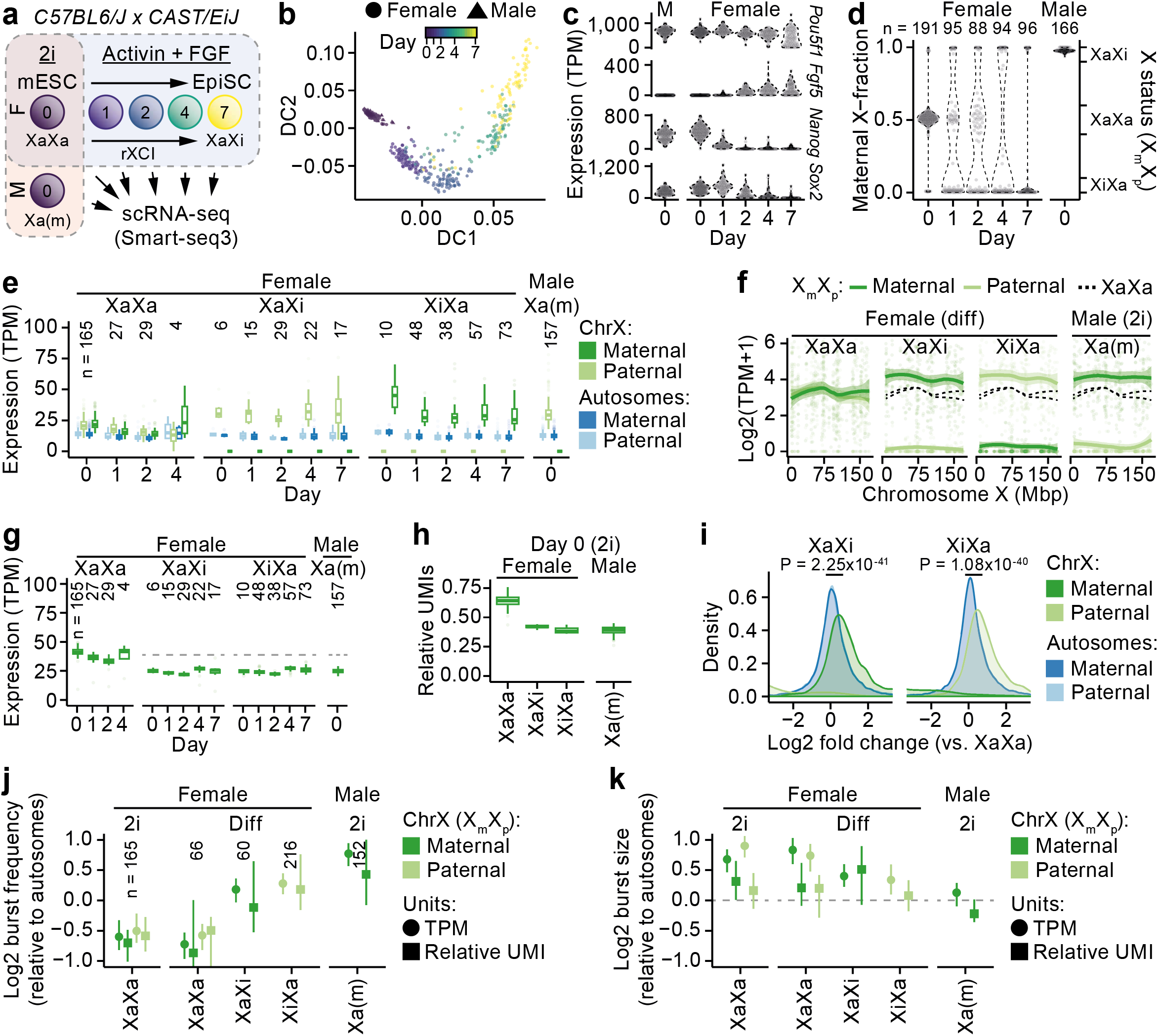
X-upregulation during X-inactivation establishment *in vitro*. **a**, Schematic overview of experimental setup. **b,** Diffusion map of mESC differentiation for 687 cells sequenced using Smart-seq3. **c-d**, Expression of pluripotency and differentiation marker genes *Pou5f1, Fgf5, Nanog* and *Sox2* (c) and maternal fraction of X-chromosome expression (d) along differentiation time. **e**, Allelic expression across differentiation days grouped by rXCI status for autosomal (n = 15,683–16,543) and X (n = 508–518) genes. **f**, Smoothed mean (LOESS fit ± 95% confidence interval) of allelic expression across the X chromosome (n = 508–524 genes). **g**, Same as (E) but for global expression for X genes (n = 1,158–1,337). Dashed line indicates XaXa median. **h**, Relative UMI expression for mESCs under 2i conditions for X genes (n = 805). **i**, Density plots of gene-wise fold changes of matching alleles between XaXa and XCI state (XaXi or XiXa; C57 and CAST allele active, respectively) in female differentiating mESCs for autosomal (n = 13,696–14,845) and X (n = 383–433) genes. Mann-Whitney *U* test P-values for active alleles. **j-k**, Relative burst-frequency (j) and size (k) for active X alleles (Xa) grouped by growth condition and rXCI status. Data shown as median ± 95% confidence interval for TPM and relative UMIs for autosomal (n = 10,836–16,714) and X (n = 348–545) genes.

Previous studies have linked XCU to increased transcriptional initiation (*3*, *13*, *14*) and bursting (*15*), suggesting that XCU is driven by transcriptional events on the active allele. We inferred parameters of transcriptional burst kinetics at the resolution of single alleles using both molecule- (UMI) and read-count (TPM) data (**Methods**), revealing moderate and balanced burst frequencies for the two XaXa alleles whereas all states of a singly active X allele displayed markedly increased burst frequency (FDR_TPM_ ≤ 3.97 × 10^−10^, FDR_UMI_ ≤ 5.68 × 10^−03^, FDR corrected Mann-Whitney *U* test; **Fig. 1j**). Burst sizes were not increased but remained largely constant (FDR ≥ 0.08, **Fig. 1k**), consistent with our previous findings that XCU is driven by increased transcriptional burst frequencies (*15*).

### Two waves of X-upregulation during early mouse development

Naïve mESCs are derived from, and mimic, the inner cell mass (ICM) of preimplantation stage embryos. To obtain an allelic map of XCU dynamics throughout the *in vivo* pre- and peri-implantation development, we leveraged the large allele-level scRNA-seq datasets we recently generated for the C57BL/6J *x* CAST/EiJ F1 breed across early murine embryogenesis (*16*–*18*), the developmental window encompassing the main events of XCI (*1*); imprinted XCI (iXCI), X-chromosome reactivation (XCR) and rXCI (**Fig. 2a**). We calculated X-chromosome expression at cell and allelic resolution and observed initiation of XCU at the 4-cell stage in males, *i.e*. around completion of zygotic genome activation (ZGA) (**Fig. 2b**), whereas female cells displayed balanced allelic expression until the 8-to-16-cell stage (**Fig. 2b**) coinciding with iXCI. This implies that XCU is initiated in concert with unbalanced chromosomal dosage in a sex-specific manner. We observed maintained XCU throughout preimplantation blastocyst development in both sexes (**Fig. 2b**). To investigate XCU in the emerging lineages in this window of development, we first performed graph-based clustering of blastocyst cells into ICM, trophectoderm (TE) and two immature clusters, followed by hierarchical clustering of ICM into epiblast (EPI) and primitive endoderm (PrE) based on known lineage markers (**Supplementary Fig. 2a-b, Methods**). This showed XCU to be present in all lineages of the late blastocyst, including EPI cells prior to XCR (**Supplementary Fig. 2c**), again suggesting that XCU is regulated based on active X alleles. Indeed, during post-implantation development (E5.5-6.5), male cells (independent of lineage) and female cells of extraembryonic lineages (visceral endoderm, VE; extraembryonic ectoderm, ExE) retained XCU along with iXCI across all timepoints (**Fig. 2c**). As expected, female E5.5 EPI cells, residing in reactivated XaXa state, showed X-linked expression close to autosomal levels for each allele (**Fig. 2c**), importantly demonstrating erasure of XCU *in vivo*. This was followed by a second wave of XCU in cells where rXCI was either underway or completed (E6.0–6.5) (**Fig. 2c**). As rXCI is an asynchronous process (*16*, *17*), we stratified and analyzed EPI cells based on rXCI status which verified that cells of XaXa state expressed each X allele at moderate levels, similar to autosomes (**Fig. 2d**), consistent with our *in vitro* findings (**Fig. 1**). Intriguingly, XCU displayed gradual chromosome-wide re-establishment along with rXCI establishment (**Fig. 2d-e**), showing that XCU is not a trigger-like mechanism but a dynamic ‘tuning-like’ process. To control for effects related to differentiation, we inferred pseudotime differentiation trajectories for each lineage (**Supplementary Fig. 2d-e**, **Methods**), demonstrating minimal association with X-linked expression compared to allele usage (**Supplementary Fig. 2f**).

**Figure 2.**
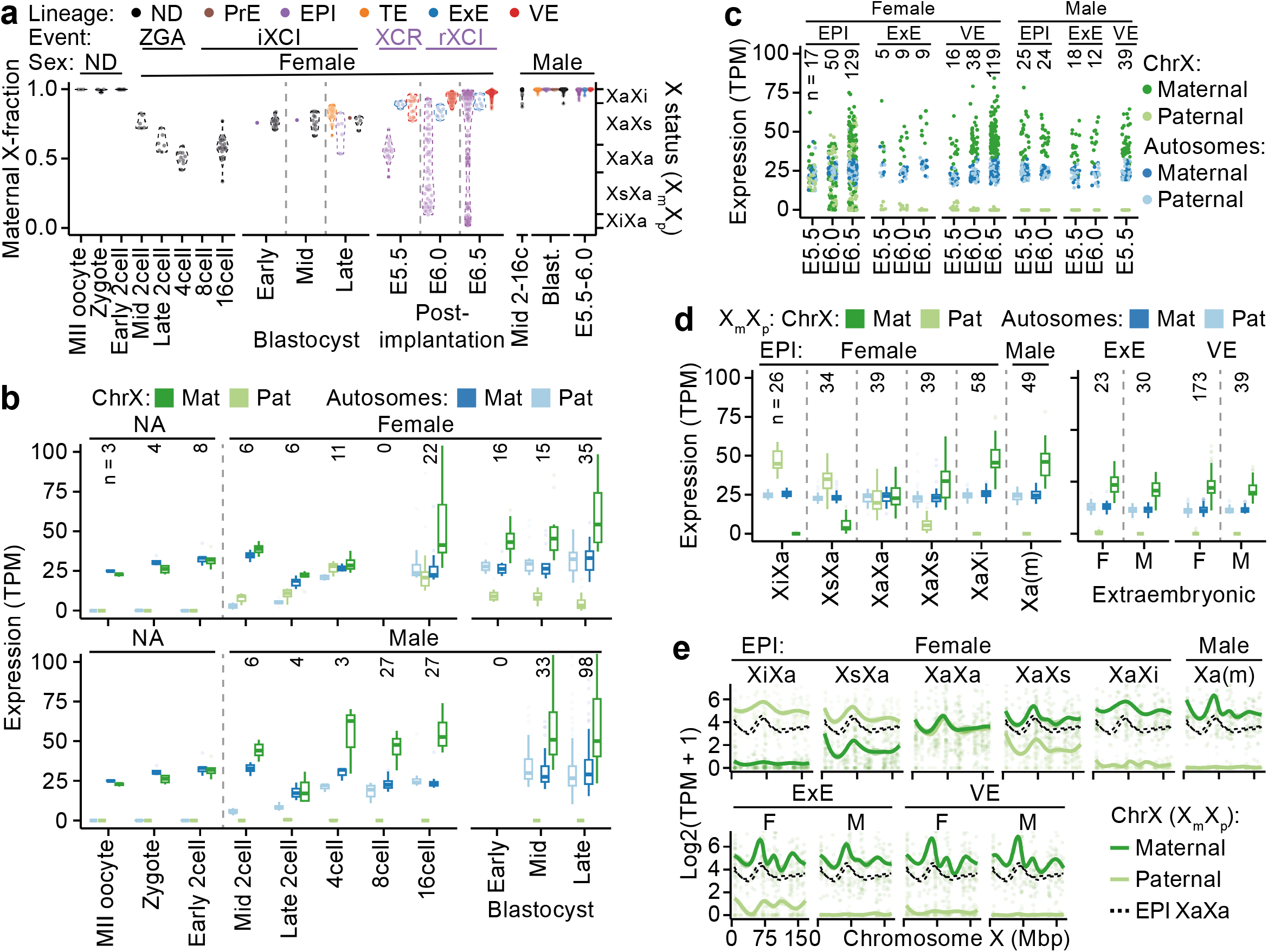
The dynamics of X-upregulation *in vivo*. **a**, Maternal fraction X-chromosome expression across female mouse early embryo development with key events of X-chromosome regulation labelled above; ZGA (mid-2- to 4-cell stage), iXCI (initiating at 8- to 16-cell stage), XCR (epiblast-specific, late blastocyst to early implantation) and rXCI (epiblast-specific, following XCR). Male data shown to the right. ND = Sex not determined. **b**, Expression levels per allele of pre-implantation development for autosomal (n = 11,164–13,347) and X (n = 323–437) genes. **c-d**, Allelic expression of post-implantation embryos grouped by embryonic day (c) or rXCI status (d) for autosomal (n = 14,941–17,643) and X (n = 597–713). **e**, Smoothed mean (GAM fit ± 95% confidence interval) of allelic expression across the X chromosome (genes n = 543–659), dashed lines represent EPI XaXa expression.

### A linear relation of X-chromosome upregulation with X-inactivation

If allelic X-chromosome expression dosage is compensated based on output rather than the number of active chromosome copies, such a mechanism would ensure similar mRNA levels regardless of degree of XCI. Indeed, allelic X-linked expression was highly correlated with maternal fractions in EPI cells (adjusted R^2^_maternal_ = 0.80, P_maternal_ = 2.92 × 10^−69^, adjusted R^2^_paternal_ = 0.91, P_paternal_ = 1.19 × 10^−103^, linear regression; **Fig. 3a and Supplementary Fig. 3a**). Surprisingly, genes escaping XCI showed similar trends to other X-linked genes (P_maternal_ = 0.62, P_paternal_ = 0.71, Kolmogorov–Smirnov test; **Supplementary Fig. 3a**) which would be unlikely if XCU is independently regulated per gene and allele. Furthermore, variability of iXCI completeness in extraembryonic lineages and pre-implantation stages also followed the trend projected from EPI cells (**Fig. 3a and Supplementary Fig. 3b**), suggesting that the first, iXCI-associated, wave of XCU is achieved by the same mechanism.

**Figure 3.**
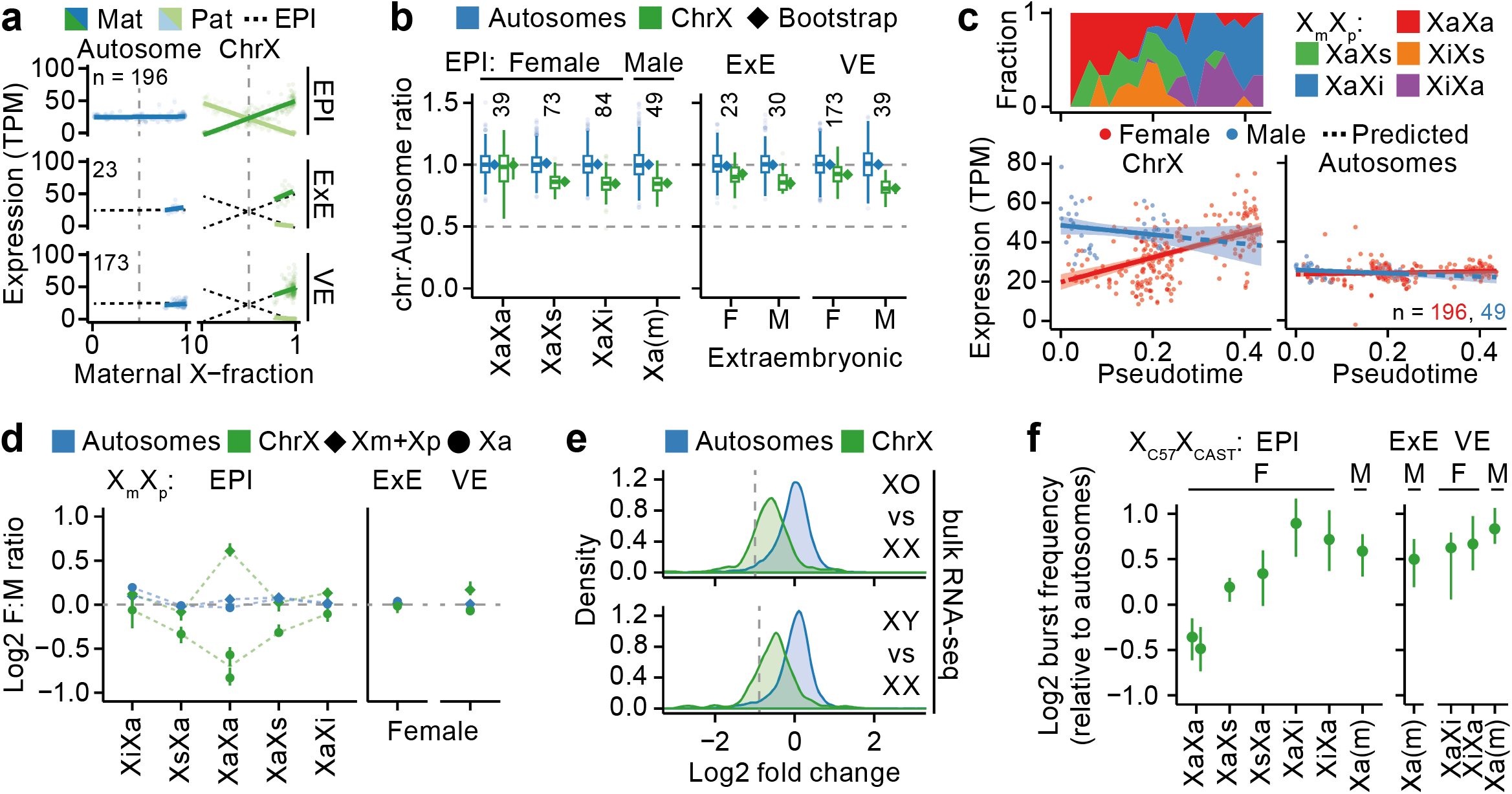
Linear relationship between X-upregulation and X-inactivation. **a**, Scatter of expression level per allele and maternal X-fraction per female cell and lines indicating linear model mean ± 95% confidence interval, shown for three lineages. Dashed black lines projects EPI and dashed gray lines indicate maternal fraction 0.5. **b**, Chromosomal expression ratios shown as box plots or bootstrapped medians (♦) ± 95% confidence interval. Expression-level threshold 10 TPM. F = female, M = male. **c**, Binned X status (top) and expression of Xa allele shown as linear model mean ± 95% confidence interval for autosomal (n = 17,643) and X (n = 713) genes (bottom) along pseudotime trajectory in EPI cells. Dashed lines represent values predicted from linear model. **d**, Female:male ratios for the active X allele (Xa; •) or global expression (Xm+Xp; ♦) shown as median and ± 95% confidence interval. **e**, Density plot of differential expression in mESC bulk RNA-seq data (XO n = 2, XX n = 12, XY n = 14) for autosomal (n = 10,961) and X (n = 405) genes. Dashed line indicates expected half of autosomal expression. **f**, Burst frequencies of the Xa allele (n = 252–V316 genes) relative to autosomal alleles of same strain (n = 9,204–V10,577 genes), shown as median and 95% confidence interval. **a-b**, **d**, Autosomal (n = 14,941–V17,643) and X (n = 597–V713) genes.

Notably, previous studies lacking allelic resolution approximated XCU using X:autosome (X:A) expression ratios (*3*–*6*) for which the tuning-like compensation (<u>Xa</u>Xi > <u>Xa</u>Xs > <u>Xa</u>Xa, considering expression of one active allele in each state) would be undetectable. Indeed, X:A ratios were indistinguishable in non-XaXa cells, regardless of sex, lineage and rXCI status (median EPI = 0.85–0.86 (XaXa = 0.98); VE = 0.81–0.93; ExE = 0.86–0.90; **Fig. 3c and Supplementary Fig. 3c-d, Methods**). Interestingly, ancestral X-Y gene pairs were not upregulated in males whereas corresponding X-linked gametologs were subject to XCU in females (**Supplementary Fig. 3e-g and Supplementary Table 2**).

Due to caveats with indirect comparisons across chromosomes (*i.e.* X:A ratios), we sought to directly assess the dosage balancing dynamics of XCU. As male cells persistently exhibited XCU past ZGA, we compared male and female Xa alleles along rXCI which indeed revealed that expression of the female Xa allele gradually increased towards male X levels as rXCI progressed (**Fig. 3c & Supplementary Fig. 3h**). We extended this comparison by calculating gene-wise female:male expression ratios at allelic- (Xa) and global level (Xm+Xp; sum of maternal and paternal allele) per lineage, embryonic day and rXCI state (**Methods**). This showed that global gene-level expression was balanced in all except XaXa cells whereas female Xa alleles showed progressive upregulation along rXCI (**Fig. 3d**), consistent with balanced mRNA levels between both alleles. The increased relative expression in XaXa cells is in agreement with our *in vitro* results (**Fig. 1h-i**) and was further supported by reanalysis of bulk RNA-seq data (**Methods**) where mESCs of male sex (XY) and female cells lacking one X-chromosome copy (XO) displayed lower X-linked expression compared to XX cells (**Fig. 3e**), confirming our *in vitro* observation that XCU of one allele does not fully reach the combined levels of both alleles in cells of XaXa state.

If XCU tunes transcription based on mRNA levels, we expect transcription from the Xa allele to be merely slightly increased in cells with partially established XCI due to residual mRNAs from the other allele. We inferred transcriptional kinetics of XCU *in vivo* and indeed found that burst frequencies on the Xa allele were progressively increased during rXCI establishment (**Fig. 3f**) whereas burst sizes remained similar to autosomal levels upon XCU (**Supplementary Fig. 3i**). We furthermore inferred transcriptional kinetics in extraembryonic cells subject to iXCI, also revealing increased burst frequencies for the Xa allele (**Fig. 3f**), again pointing to a unified XCU mechanism in all lineages.

## Discussion

By allele-level mRNA measurements in single cells throughout embryonic stem cell differentiation *in vitro* and early development *in vivo*, we have provided a unified model of XCU and its interplay with the XCI dynamics, schematically summarized in **Fig. 4**. We show distinct sex-specific initiation of XCU where X:autosome dosage would otherwise be unbalanced; after completion of ZGA in males and together with iXCI in females (**Fig. 2b**). This is in contrast to previous studies suggesting XCU to be established already in the zygote (*6*) or progressively after the 4-cell stage in both sexes (*7*, *8*) using X:A ratios. These discrepancies may be explained by the inability of non-allelic measurements to correctly distinguish XCU in the presence of other allelic processes such as ZGA and XCI, whereas we in this study directly attribute XCU to the active X allele. Furthermore, we reveal that naïve mESCs cells lack XCU whereas previous studies suggested XCU to be present in these cells (*3*, *5*, *19*), which could be explained by the fact that total XaXa expression levels are higher than what is achieved through XCU (**Fig. 1g–i and Fig. 3b, d–e**).

What is the mechanism underlying the XCU dynamics? Our findings are in line with a recently proposed model (*20*) in which expression balance is achieved by local transcription factor concentrations. As XCI is gradually established (*21*), transcription factors would be available and shift to the active X allele, increasing transcriptional burst frequencies (*22*) resulting in increased mRNA levels. This would explain why active histone modifications do not scale proportionally with mRNA levels on the hyperactive X allele (*13*, *14*) and why genes that escape XCI are subject to similar dosage balancing as other X-linked genes (**Supplementary Fig. 3e–g**). Enhancer elements likely play key roles in mediating XCU as they may not only increase local transcription factor concentrations through DNA contacts (*23*) but also regulate transcriptional burst frequencies (*24*) which is a driver of XCU (*15*).

**Figure 4.**
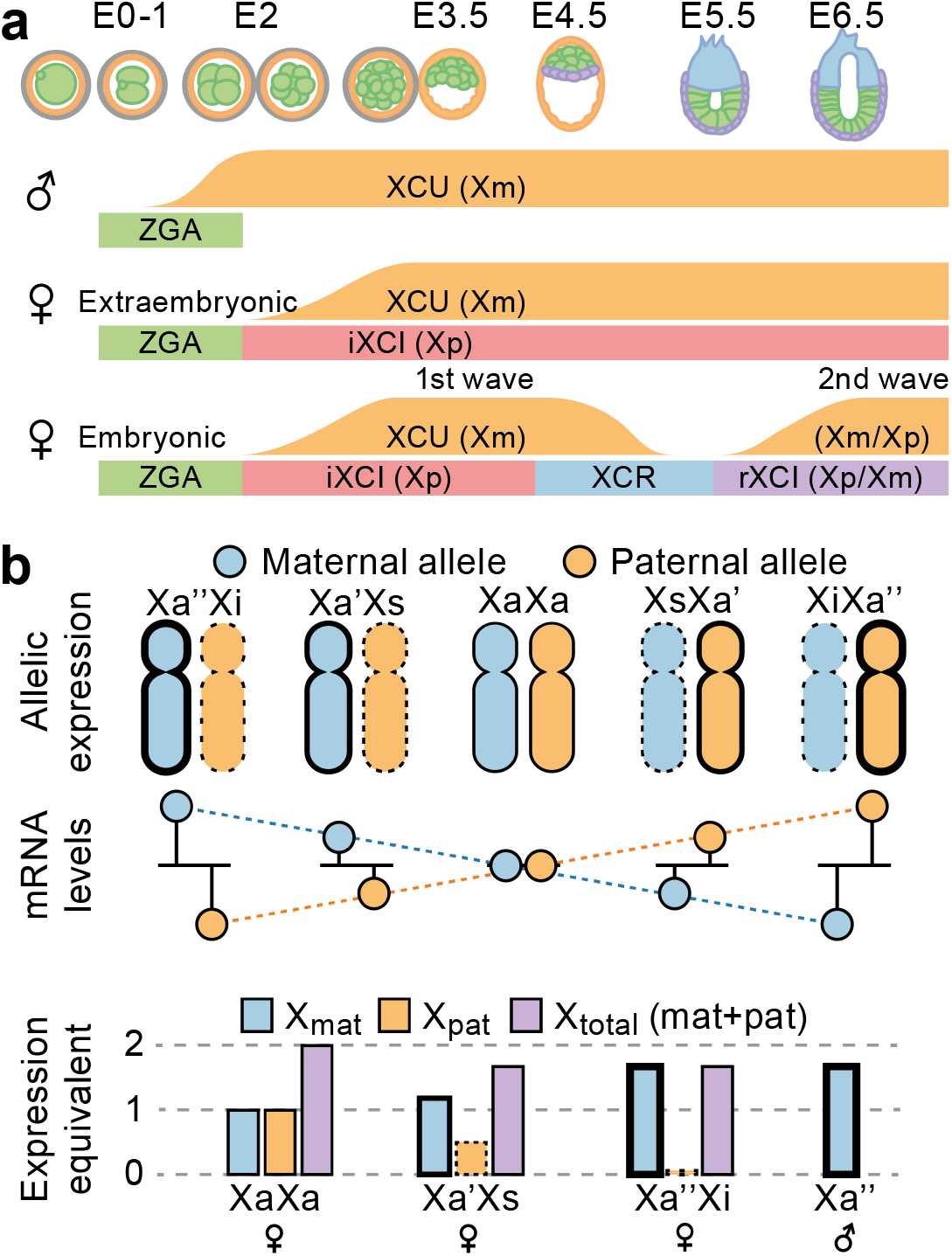
Model of X-upregulation. **a**, Schematic of sex- and lineage-specific dynamics of X-upregulation (XCU) across mouse embryonic development. **b**, Proposed model of XCU as a flexible mechanism of allelic dosage balancing. As one allele is gradually silenced, the concentration of available transcription factors is shifted towards the active allele (Xa), increasing transcriptional burst frequencies and mRNA levels in a linear fashion (top). However, the single hyperactive allele (Xa”) does not fully reach the combined expression level of two moderately active alleles in XaXa state (bottom).

Our findings that XaXa cells lack XCU (**Fig. 1g-h and Fig. 2c-d**) is in line with observations of balanced X:A dosage in haploid cells (including MII oocytes; **Fig. 2b**) and cells with two active X chromosomes, such as primary oocytes and primordial germ cells (*6*, *25*–*27*). Two active X chromosomes is considered a hallmark of the naïve female stem cell state and a gold standard in reprogramming studies (*28*). Our model of two moderately active alleles in the XaXa state in contrast to hyperactive female XaXi and male XaY alters the interpretation of X-chromosome expression level measurements for assessment of reprogramming success and naïveness. Furthermore, in light of our current results in mouse, it is plausible that the “dampened” biallelic (XaXa) X-chromosome state reported in female human preimplantation development (*29*–*31*) and human naïve pluripotent stem cells (*32*) reflect lack or erasure (*33*) of XCU.

Together, our study provides a unified model of XCU and its link to XCI, and evidence for flexible balancing of allelic expression levels to compensate dosage of the X chromosome.

## Acknowledgements

This study was made possible by grants to B.R. from the Swedish Research Council (2017-01723), Åke Wiberg’s Foundation, and the Ragnar Söderberg Foundation. We thank members of the Reinius lab for discussions and comments on the manuscript.

## Contributions

A.L.: Investigation, Methodology, Formal analysis, Conceptualization, Visualization, Data Curation, Writing - Original Draft, Writing - Review & Editing. C.C.: Investigation. N.A.: Investigation. M.E.: Resources. Q.D.: Methodology, Resources. B.R.: Investigation, Methodology, Conceptualization, Supervision, Resources, Project administration, Funding acquisition, Writing - Original Draft, Writing - Review & Editing.

## Data and materials availability

Raw and pre-processed Smart-seq3 data generated is publicly available at ArrayExpress under accession E-MTAB-9324. Previously published raw data is available at Gene Expression Omnibus under accessions GSE45719, GSE74155, GSE109071, GSE23943 and GSE90516. Data and code generated during this study are available at github: github.com/reiniuslab/Lentini_XCU_in_vivo.

## Materials and Methods

### Ethics statement

All animal experimental procedures were performed in accordance with Karolinska Institutet’s guidelines and approved by the Swedish Board of Agriculture (permits N225/15, N343/12 and 18729-2019 Jordbruksverket).

### Derivation and culturing of cell lines

Male and female cell lines were established as previously described (*16*). In brief, mESCs were derived from E4 blastocysts of F1 embryos (female C57BL/6J × male CAST/EiJ) and adapted to 2i condition by growing them in gelatin-coated flasks in N2B27 medium (50% neurobasal medium [Gibco], 50% DMEM/F12 [Gibco], 2 mM L-glutamine [Gibco], 0.1 mM β-mercaptoethanol, NDiff Neuro-2 supplement [Millipore], B27 serum-free supplement [Gibco]) supplemented with 1,000 units/mL LIF, 3 μM Gsk3 inhibitor CT-99021, 1 μM MEK inhibitor PD0325901, and passaged with accutase [Gibco]. To induce differentiation toward EpiSCs, mESCs grown in serum/LIF were plated on FBS coated tissue culture plates (coated overnight at 37°C) in N2B27 medium supplemented with 8 ng/mL Fgf2 (R&D) and 20 ng/mL Activin A (R&D) at a cell density of 1 × 10^4^ cells/cm^2^ and cultured for up to 7 days. Cells were split and re-plated in the same condition after 1, 2, 4 and 7 days of differentiation. At each split an aliquot of cells was collected for single-cell sorting into 96-well plates containing Smart-seq3 lysis buffer.

### Single-cell RNA sequencing (Smart-seq3)

scRNA-seq libraries were constructed as previously described (*10*) with slight modification. Briefly, cells were single-cell sorted into 96-well low-bind PCR-plates [Eppendorf] containing 3 μl of lysis buffer (0.5 units/μl RNase inhibitor [Takara], 0.15% Triton X-100 [Sigma], 0.5 mM (each) dNTP [Thermo Scientific], 1 μM oligo-dT primer [5′-biotin-ACGAGCATCAGCAGCATACGAT30VN-3′; IDT], 5% PEG [Sigma]). Sorting was performed using an SH800 [Sony]. Plates were briefly centrifuged immediately after sorting, sealed, and stored at −80°C. For cell lysis and RNA denaturation, plates were incubated at 72°C for 10 min and immediately placed on ice. Next, 5 μl of reverse transcription mix (50 mM Tris-HCl, pH 8.3 [Sigma], 75 mM NaCl [Ambion], 1 mM GTP [Thermo Scientific], 3 mM MgCl_2_ [Ambion], 10 mM DTT [Thermo Scientific], 1 units/μl RNase inhibitor [Takara], 2 μM of template-switch oligo [5′-biotin-AGAGACAGATTGCGCAATGNNNNNNNNrGrGrG-3′; IDT] and 2 U/μl of Maxima H-minus reverse transcriptase [Thermo Scientific]) was added to each sample. Reverse transcription was carried out at 42°C for 90 min followed by 10 cycles of 50°C for 2 min and 42°C for 2 min and the reaction was terminated at 85°C for 5 min. PCR pre-amplification was performed directly after reverse transcription by adding 17 μl of PCR mix (containing DNA polymerase, forward and reverse primer) bringing the final concentration in the 25 μl reaction to 1x KAPA HiFi ReadyMix [Roche] 0.1 μM forward primer [5′-TCGTCGGCAGCGTCAGATGTGTATAAGAGACAGATTGCGCAATG-3′; IDT] and 0.1 μM reverse primer [5′-ACGAGCATCAGCAGCATACGA-3′; IDT]). Thermocycling was performed as follows: 3 min at 98°C, 22 cycles of 20 s at 98°C, 30 s at 65°C and 6 min at 72°C, and final elongation at 6 min at 72°C. After PCR pre-amplification, samples were purified with AMpure XP beads [Beckman Coulter] at volume ratio 0.8:1. Library size distributions were monitored using high-sensitivity DNA chips (Agilent Bioanalyzer 2100) and cDNA concentrations were quantified using the Quant-iT PicoGreen dsDNA Assay Kit [Thermo Scientific]. cDNA was subsequently diluted to 100–200 pg/μl.

Tagmentation was performed using in-house produced Tn5 (*34*). 2 ng of cDNA in 5 μl water was mixed with 15 μl tagmentation mix (0.2 μl Tn5, 2 μl 10x TAPS MgCl_2_ Tagmentation buffer; 5 μl 40% PEG-8000; 7.8 μl water, per reaction) and incubated 8 min at 55°C in a thermal cycler. Tn5 was inactivated and released from the DNA by the addition of 4 μl 0.2% SDS and 5 min incubation at room temperature. Library amplification was performed by adding 5 μl mix of 1 μM of forward and reverse custom-designed Nextera index primers [forward: 5′-CAAGCAGAAGACGGCATACGAGATNNNNNNNNNNGTCTCGTGGGCTCGG-3′, reverse: 5′-AATGATACGGCGACCACCGAGATCTACACNNNNNNNNNNTCGTCGGCAGCGTCIDT-3′, where N represents the 10-bp index bases; IDT] and 15μl PCR mix (1μl KAPA HiFi DNA polymerase [Roche]; 10μl 5x KAPA HiFi buffer; 1.5 μl 10 mM dNTPs; 3.5μl water, per reaction), and thermal cycling: 3 min 72°C, 30 s 95°C, 13 cycles of 10 s 95°C; 30 s 55°C; 30 s 72°C, followed by final elongation at 5 min 72°C; 4°C. DNA sequencing libraries were purified using 0.8:1 volume of AMPure XP beads [Beckman Coulter]. Libraries were sequenced using a NextSeq 550 and High Output kits [Illumina].

### Smart-seq3 data analysis

A hybrid mouse genome index was constructed by N-masking the reference genome (GRCm38_68) for CAST/EiJ SNPs from the Mouse Genomes Project (mgp.v5.merged.snps_all.dbSNP142) (*35*) using SNPsplit (0.3.2) (*36*). Raw Smart-seq3 data was processed using zUMIs (2.8.0) (*37*). Briefly, sample barcodes were filtered and data was aligned with STAR (2.7.2a) (*38*) (options: --clip3pAdapterSeq CTGTCTCTTATACACATCT --quantMode TranscriptomeSAM) and reads were assigned to both intron and exon features (Mus_musculus.GRCm38.97.chr.gtf) using FeatureCounts (*39*). Next, barcodes were collapsed using 1 hamming distance and gene expression was calculated for both reads and UMIs. Finally, allele-level expression was calculated from the zUMIs output as previously described (https://github.com/sandberg-lab/Smart-seq3/tree/master/allele_level_expression) (*10*). 77 cells were excluded due to low read depth (>3 MADs).

### Processing of published C57BL/6J × CAST/EiJ scRNA-seq data

Pre-processed Smart-seq data for MII oocyte – 16-cell stages were obtained (*18*) and two cells were excluded (8cell_8-3 & 16cell_4-2) due to potential sample problems (high paternal X ratio in male and non-ZGA with clustering together with Zygote samples, respectively). Pre-processed blastocyst Smart-seq2 data was obtained (*18*) and 18 late blastocysts were excluded due to low read depth (>3 MADs). Pre-processed Smart-seq2 data for post-implantation embryos was obtained (*16*, *17*).

### Genes escaping X inactivation

A list of known mouse escapee genes (*1810030O07Rik*, *2010000l03Rik*, *2010308F09Rik*, *2610029G23Rik*, *5530601H04Rik*, *6720401G13Rik*, *Abcd1*, *Araf*, *Atp6ap2*, *BC022960*, *Bgn*, *Car5b*, *D330035K16Rik*, *D930009K15Rik*, *Ddx3x*, *Eif1ax*, *Eif2s3x*, *Fam50a*, *Flna*, *Ftsj1*, *Fundc1*, *Gdi1*, *Gemin8*, *Gpkow*, *Huwe1*, *Idh3g*, *Igbp1*, *Ikbkg*, *Kdm5c*, *Kdm6a*, *Lamp2*, *Maged1*, *Mbtps2*, *Med14*, *Mid1*, *Mmgt1*, *Mpp1*, *Msl3*, *Ndufb11*, *Nkap*, *Ogt*, *Pbdc1*, *Pdha1*, *Prdx4*, *Rbm3*, *Renbp*, *Sh3bgrl*, *Shroom4*, *Sms*, *Suv39h1*, *Syap1*, *Tbc1d25*, *Timp1*, *Trap1a*, *Uba1*, *Usp9x*, *Utp14a*, *Uxt*, *Xist*, *Yipf6*) was compiled from previous work (*7*, *40*–*42*) and excluded from analyses of positional expression levels and expression ratios (described below).

### Expression calculations

Allelic reference ratios were calculated as Counts_C57_/Counts_Total_ after exclusion of the top 10% expressed X-linked genes to avoid bias from highly expressed X genes. XCI status was determined as active (Xa/Xa), semi-inactivated (Xa/Xs) or fully inactivated (Xa/Xi) for allelic ratios in the intervals (0.4, 0.6), [0.6, 0.9) and [0.9, ∞) and the inverse, respectively. TPM was calculated for gene *i* in cell *j* as TPM_ij_ = (FPKM_i_ / ∑_j_ FPKM_j_) × 10^6^ and allelic TPM was calculated by scaling TPM by reference ratios per gene and cell. Relative UMIs was calculated like TPM but using UMI counts and an offset of 10^2^. A gene was considered expressed in a dataset/lineage if the average TPM expression was > 0. To obtain a robust estimate for whole-chromosome expression, 20% trimmed TPM means was calculated per cell.

### Expression ratios

X:Autosomal ratios are well-known to be highly dependent on the thresholds used (*3*, *11*, *14*) due to X-chromosome-specific expression patterns (**fig. S3C**), therefore we tested several different thresholds using XaXa cells (**fig. S3D**). Two different threshold were selected; >10 TPM as the X expression distribution was comparable to autosomal levels in XaXa cells (see **fig. S3C**, used in **Fig. 3B**), and >1 TPM to be comparable to previous studies (used in **fig. S3E & G**) (*3*, *5*, *14*). Chromosome:autosome expression ratios were calculated as TPM relative to median of autosomes after applying expression thresholds and excluding genes that escape XCI. To account for different number of genes between chromosomes, a bootstrapping method was used (*7*). For bootstrapped ratios, random autosomal gene sets of the same size were selected as a background, repeated n = 10^3^ times.

Female:male ratios were calculated after exclusion of X escapees as TPM relative to gene average per embryonic day and lineage, as well as for the active X allele for allelic data.

### Differential expression of scRNA-seq data

For Smart-seq3 data, global count data was size-factor normalized using scater (1.12.2)/scran (1.12.1) (*43*) and genes expressed in >10% of cells were kept. Differential expression was calculated along days of differentiation as a continuous variable using likelihood ratio tests as implemented in MAST (1.10.0) (*44*). Gene set enrichment for mouse GO biological process gene sets obtained from MGI (http://www.informatics.jax.org/downloads/reports/index.html#go; accessed 2020-06-23) was performed on the differential expression model against a bootstrapped (n = 100) control model as implemented in MAST.

### Dimensionality reduction, clustering and trajectory inference

For Smart-seq3 data, highly variable genes (HVGs) were identified from size-factor-normalized counts using scater/scran and ordered by biological variance and FDR. Data was visualized for top 1,000 HVGs using diffusion maps (*45*).

For *in vivo* blastocyst data, HVGs were obtained as explained above and dimensionality reduction was performed for top 1,000 HVGs using UMAP (*46*) and cells were Louvain clustered based on top 1,000 HVG ranks using scran.

For *in vivo* post-implantation data, pseudotime trajectories were inferred using Slingshot (1.2.0) (*47*) from normalized counts following Mclust (5.4.5) (*48*) clustering on diffusion map coordinates.

### X-Y homology

List of X-Y homologs was obtained from (*49*) and sequence similarity of expressed X-Y homolog transcripts was calculated using the nucleotide BLAST webservice (*50*) using default settings (performed 2020-03-24).

### Kinetics inference

Missing allelic data points were set to 0 if the gene was detected on the other allele. Kinetic parameters were calculated per lineage/genotype/XCI status/growth condition (depending on dataset) using txburst (https://github.com/sandberg-lab/txburst) (*24*). Genes not passing filtering steps were excluded and relative burst frequency (k_on_) and burst size (k_syn_/k_off_) was calculated relative to median of expressed autosomal genes (per lineage). Only groups with > 20 cells were kept to increase the reliability of the statistical inference.

### Re-analysis of published bulk RNA-seq data

Raw RNA-seq data including both male and female mESCs was obtained (*51*, *52*) and quantified to protein-coding transcripts from GENCODE vM22 (*53*) using pseudoalignment with Salmon (0.14.1) (*54*). Transcript abundance estimates were summarized to gene-level using tximport (1.12.3) (*55*) and differential expression was calculated using likelihood ratio tests in DESeq2 (1.24.0) (*56*). A full model including cellular genotype (XX, XY or XO), cell culture condition (2i or serum) and study accession was tested against a reduced model without the genotype term. Lowly expressed genes (<100 average normalized counts) were excluded from final plots. Serum growth conditions only showed a minor effect on global X expression compared to 2i (not shown), consistent with a low degree of cells exhibiting complete XCI (*16*).

### Statistics and data visualization

All statistical tests were performed in R (3.6.1) as two-tailed unless otherwise stated. Heatmaps were visualized using ComplexHeatmap (2.0.0) (*57*) and all other plots were made using ggplot2 (3.2.1) (*58*). Box plots are presented as median, first and third quartiles, and 1.5x inter-quartile range (IQR). For median ± confidence interval plots, bootstrapped 95% confidence intervals (n = 1,000) were calculated using the percentile method (*59*) as implemented in the R boot package (1.3-23).

---

**Supplementary Figure 1.**
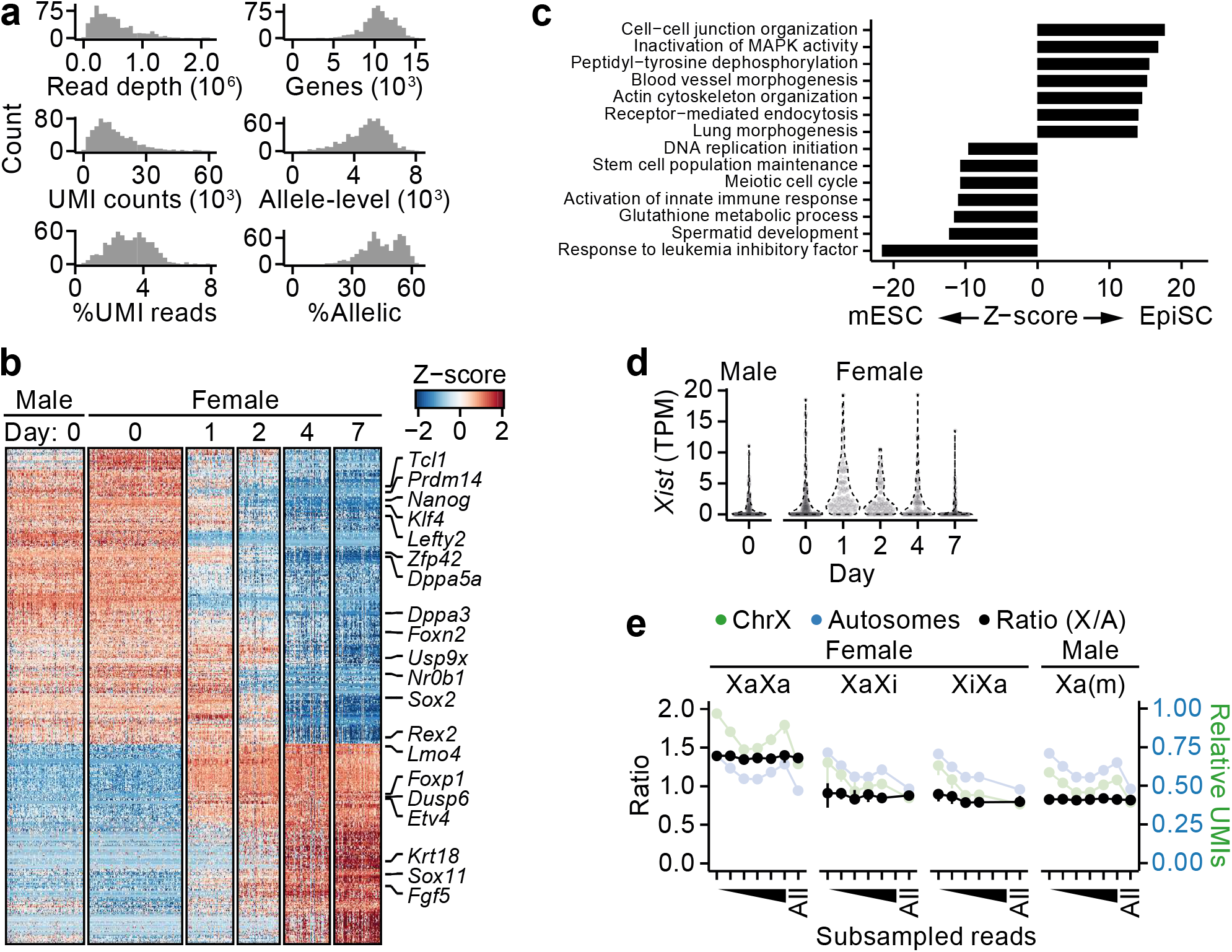
Validation of mESC differentiation. **a**, Metrics for generated Smart-seq3 data. **b**, Heatmap of differentially expressed genes along mESC differentiation. **c**, Enriched pathways for differentially expressed genes in (b). **d**, Xist expression along mESC differentiation. **e**, UMI counts for subsampled reads (n = All, 1M, 0.75M, 0.5M, 0.25M, 0.1M and 0.05M reads).

**Supplementary Figure 2.**
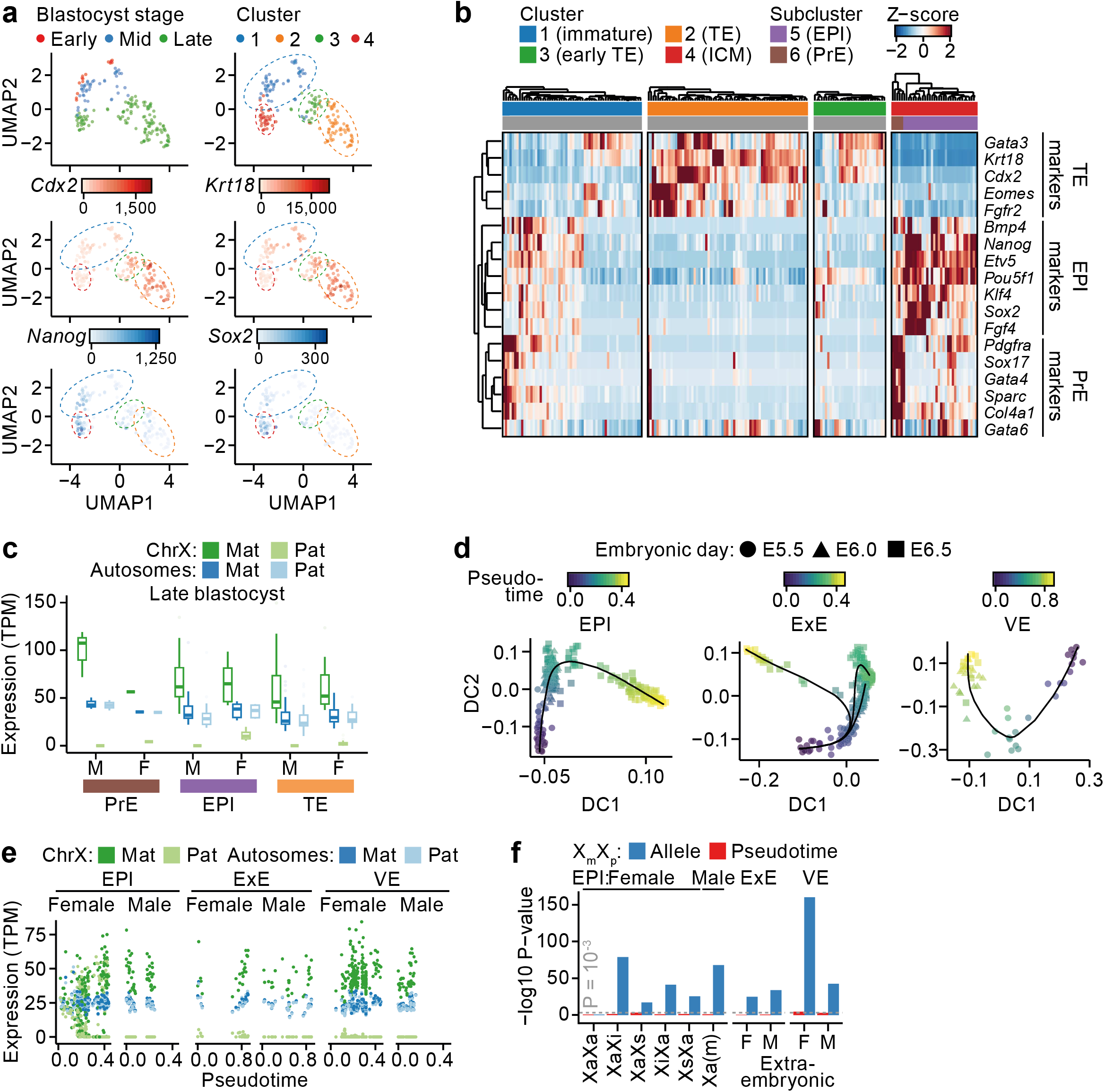
Extended analysis of early embryo development. **a**, UMAP dimensionality reduction and graph-based clustering of blastocysts. **b**, Heatmap and hierarchical clustering of blastocysts using lineage-specific marker genes. **c**, Allelic expression of late blastocyst subclusters identified in (a-b). **d**, Diffusion map dimensionality reduction and trajectory inference per post-implantation lineage. **e**, Allelic expression along pseudotime trajectories for post-implantation embryos. **f**, Association of allele usage or pseudotime on allelic expression using linear modelling.

**Supplementary Figure 3.**
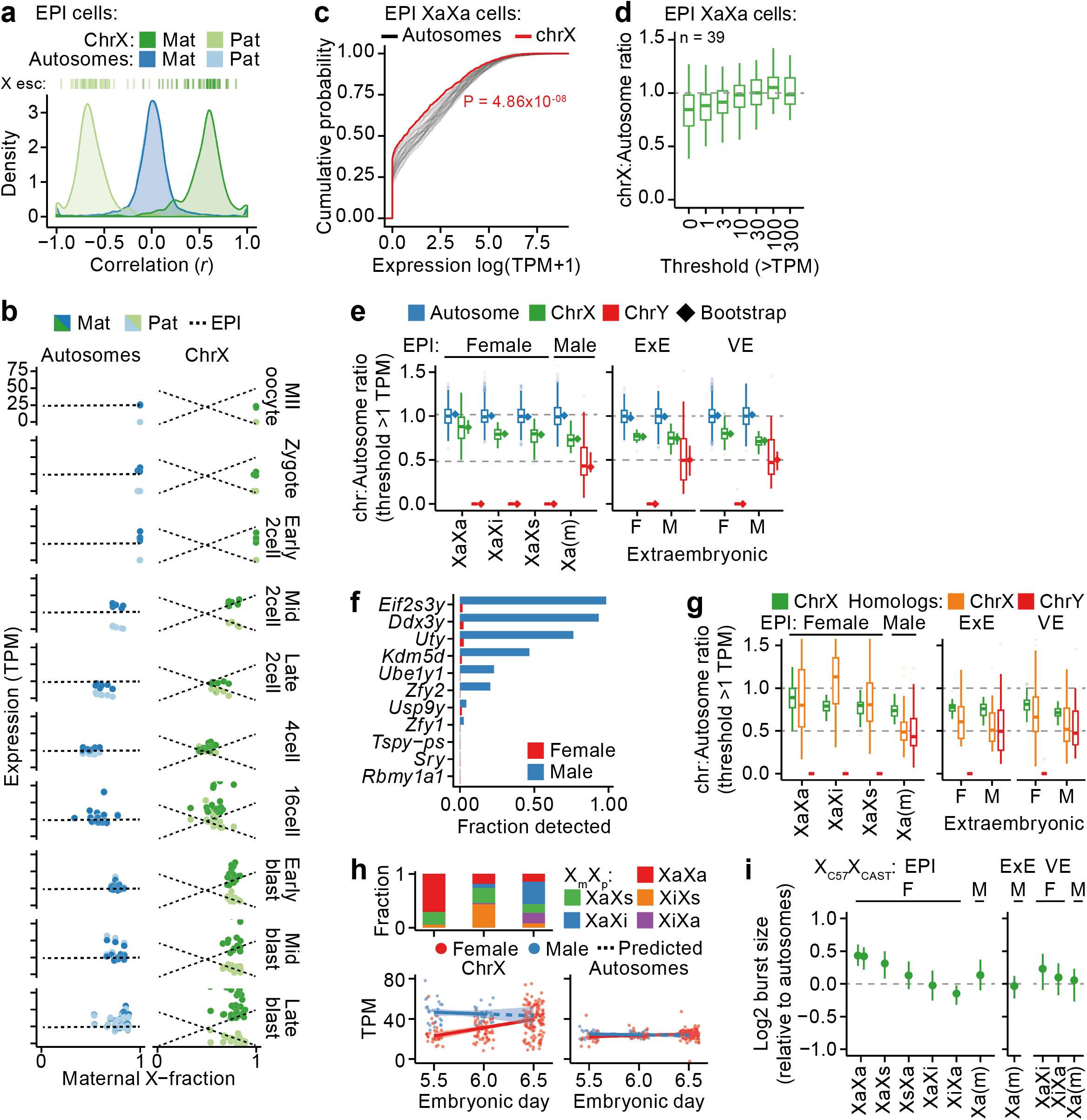
Extended analyses of linear relationship of X-upregulation. **a**, Density plot of gene-wise correlations (Pearson) against maternal X-fractions in EPI cells. X esc = X escapees. **b**, Relationship between X expression and maternal X-fraction, dashed lines represent EPI. **c**, Empirical cumulative distribution of expression for autosomes and the X chromosome. Kolmogorov-Smirnov test P-value. **d**, Effect of expression thresholds for calculating X:Autosome ratios in EPI XaXa cells. **e**, Autosomal expression ratios at >1 TPM threshold presented as box plots or bootstrapped ratios shown as median ± 95% confidence interval (♦). **f**, Fraction of expressed genes on the Y chromosome. **g**, Same as (e) but X-Y gene pair homologs shown separately. **h**, X status per time point (top). Expression of Xa allele along embryonic development day in EPI cells (bottom). Data shown as linear model mean ± 95% confidence interval, dashed lines representing values predicted from linear model. (**i**) Burst sizes of the active X allele (Xa) relative to autosomal genes, shown as median ± 95% confidence interval.

